# Fragment-based screening identifies inhibitors of the ATPase activity and of hexamer formation of Cagα from the *Helicobacter pylori* type IV secretion system

**DOI:** 10.1101/326413

**Authors:** Tarun Arya, Flore Oudouhou, Bastien Casu, Benoit Bessette, Jurgen Sygusch, Christian Baron

## Abstract

Type IV secretion systems are membrane-bound multiprotein complexes that mediate the translocation of macromolecules across the bacterial cell envelope. In *Helicobacter pylori* a type IV secretion system is encoded by the *cag* pathogenicity island that encodes 27 Cag proteins and most of these are essential for bacterial virulence. We here present our work on the identification and characterization of inhibitors of Cagα, a hexameric ATPase and member of the family of VirB11-like proteins that is essential for translocation of the CagA cytotoxin into mammalian cells. We conducted fragment-based screening using a differential scanning fluorimetry assay and identified 16 molecules that stabilize the protein during thermal denaturation suggesting that they bind Cagα. Several of these molecules affect binding of ADP and four of them inhibit the ATPase enzyme activity of Cagα. Analysis of enzyme kinetics suggests that their mode of action is non-competitive, suggesting that they do not bind to the ATPase active site. Cross-linking analysis suggests that the active molecules change the conformation of the protein and gel filtration and transmission electron microscopy show that molecule 1G2 dissociates the Cagα hexamer. Analysis by X-ray crystallography reveals that molecule 1G2 binds at the interface between Cagα subunits. Addition of the molecule 1G2 inhibits the induction of interleukin-8 production in gastric cancer cells after co-incubation with *H. pylori* suggesting that it inhibits Cagα *in vivo*. Our results reveal a novel mechanism for the inhibition of the ATPase activity of VirB11-like proteins and the identified molecules have potential for the development into antivirulence drugs.

## Author summary

We here report the results of a small-molecule screening approach to identify inhibitors of an essential virulence factor from the gastrointestinal pathogen *Helicobacter pylori*. We identifed novel chemical entities that bind to the Cagα protein and inhibit its function. We discovered a novel inhibitor binding site that disrupts the hexameric quaternary structure and inhibits ATPase enzyme activity of Cagα. Based on the structural information on the binding site, these molecules could be developed into high-affinity inhibitors that may have potential as anti-virulence drugs. This approach represents a generally applicable strategy for the inhibition of bacterial virulence factors for which structural information is available that could be applied to target similar proteins from many bacterial pathogens.

## Introduction

*Helicobacter pylori* is a widespread pathogenic bacterium that lives in the stomach of over half of the world’s population [1]. The infection with virulent strains causes inflammatory reactions, gastritis, peptic ulcers and it is one of the principal causes of stomach cancer in humans [2, 3]. Antibiotic treatments using combination therapies of three or four drugs have generally been successful, but eradication therapy is becoming increasingly difficult due to rising resistance against many antimicrobial agents, such as clarithromycin and metronidazole [4]. Novel treatment options are therefore urgently needed and targeting bacterial virulence factors to attenuate the inflammation is a strategy that could complement or even replace currently used eradication treatments.

Type IV secretion systems (T4SS) mediate the transfer of virulence factors across the cell envelope of many bacterial pathogens as well as the exchange of plasmids contributing to the spread of antibiotic resistance genes [5, 6]. *H. pylori* strains encode T4SSs that mediate the uptake of DNA as well as bacterial virulence like the *cag* pathogenicity island (*cag*-PAI)-encoded T4SS comprising 27 components of which most are essential for bacterial virulence [7–10]. The *cag*-PAI is required for the transfer of the CagA cytotoxin into mammalian cells where it is phosphorylated by Src kinase at tyrosine residues and its interactions with mammalian proteins such as SHP-2 and Grb-2 lead to rearrangements of the cytoskeleton and to proinflammatory reactions [11]. The *cag*-PAI-encoded T4SS is also a conduit for bacterial murein and for the small molecule metabolite heptulose-1,7-bisphosphate triggering signalling cascades via Nod-1 and TIFA, respectively, that contribute to the proinflammatory response [12, 13].

The *H. pylori* cag-PAI encodes 27 proteins including homologs of all 12 components of the most studied model T4SS from *Agrobacterium tumefaciens* [9]. These conserved proteins are critical for secretion system function and they are either part of surface-exposed pili, of the periplasmic T4SS core complex or they energize T4SS assembly or substrate translocation. We here focus on the Cagα (HP0525) protein that is a member of the VirB11 family of ATPases present in all T4SSs. Electron microscopic (EM) analyses and X-ray crystallography have shown that the overall structures of VirB11-like proteins from different organisms are very similar comprising homo-hexameric rings [14, 15]. The monomeric subunit consists of an N-terminal domain (NTD) and a C-terminal domain (CTD) that are linked via a short linker region comprising the nucleotide binding site. The X-ray structures of Cagα apoprotein [16], as well as of its complexes with ADP [17] and with the inhibitor ATPγS [16] have been solved. These studies revealed that the CTD forms a ‘six clawed grapple’ mounted onto the NTD, forming a hexameric ring and a dome-like chamber that is closed at one end and opened at the other [17]. Glycerol gradient centrifugation showed a large conformational change of VirB11 homologs from plasmid RP4 (TrbB) and *H. pylori* upon binding to ATP, underlining the dynamic nature of the protein [16, 18]. The other available X-ray structure from *Brucella suis* VirB11 differs from Cagα by a domain swap of the large linker region between NTD and CTD [19], but the overall structure is very similar.

Since T4SS are important for bacterial virulence they are very interesting targets for the development of drugs that disarm but do not kill bacterial pathogens [20, 21]. In our previous work, we have identified inhibitors of the dimerization of VirB8-like proteins from *B. suis* and plasmid pKM101 using the bacterial two-hybrid system and fragment-based screening approaches and we identified molecules that reduce T4SS function [22–25]. Other groups have identified peptidomimetic inhibitors of the *H. pylori* T4SS, but the targets of these molecules are not known [26]. Certain unsaturated fatty acids inhibit bacterial conjugation and the ATPase activity of the VirB11 homolog TrwD from plasmid R388, but there is no high-resolution structural information available on their binding site [27, 28]. High-throughput small molecule screening and chemical synthesis led to the identification of inhibitors of the ATPase activity of Cagα that likely bind at the ATPase active site, but structural information on their binding site is not available [29, 30]. Whereas the isolation of competitive inhibitors of the ATPase activity of VirB11 homologs is interesting, there are concerns about the specificity of these molecules since they may also inhibit other ATPases in bacteria or in mammalian cells.

To identify novel chemical entities that inhibit Cagα we here present an unbiased approach that does not specifically target its ATPase activity. To this effect, we carried out fragment based-screening using differential scanning fluorimetry (DSF) to identify molecules that bind and stabilize Cagα [31]. Four of the molecules inhibit the Cagα ATPase activity and the most potent molecule impacts the conformation of the protein and dissociates the hexamer. X-ray crystallography reveals that this molecule causes conformational changes, that it binds at the interface between Cagα monomers and it inhibits the production of interleukin-8 upon interaction between *H. pylori* and mammalian cells.

## Results

### Differential scanning fluorimetry to identify Cagα-binding fragments

To identify novel chemical entities that bind to and influence the activity of Cagα, we conducted fragment-based screening using a DSF assay. This assay measures binding of molecules to proteins by changes of the thermal melting profile in the presence of the fluorescent dye Sypro Orange [31]. We validated this assay by testing binding to previously characterized ligands that influence the conformation of Cagα, such as MgCl_2_, ADP and the non-hydrolysable substrate analog ATP-γ-S. Addition of the nucleotide ligand MgCl_2_ increases the melting temperature from 37°C to 42°C, but in the presence of MgCl_2_ and ADP or ATP-γ-S, strong increases of the melting temperature to 55°C and 60°C were observed, respectively (Fig. 1). The optimized assay conditions were used to screen a library of 505 fragments [24, 32] (supplementary Fig. 1) and 16 molecules (supplementary Fig. 2) were identified that reproducibly increase the melting temperature of Cagα by 1°C to 4°C, which is the typical range observed for binding fragments (supplementary Fig. 1). Interestingly, incubation of many of these fragments in the presence of MgCl_2_ and ADP reduces the melting temperature when compared to MgCl_2_ and ADP alone, suggesting that they impact the conformation of Cagα in a way that changes binding of the other ligands (supplementary Fig. 2).

**Figure 1.**
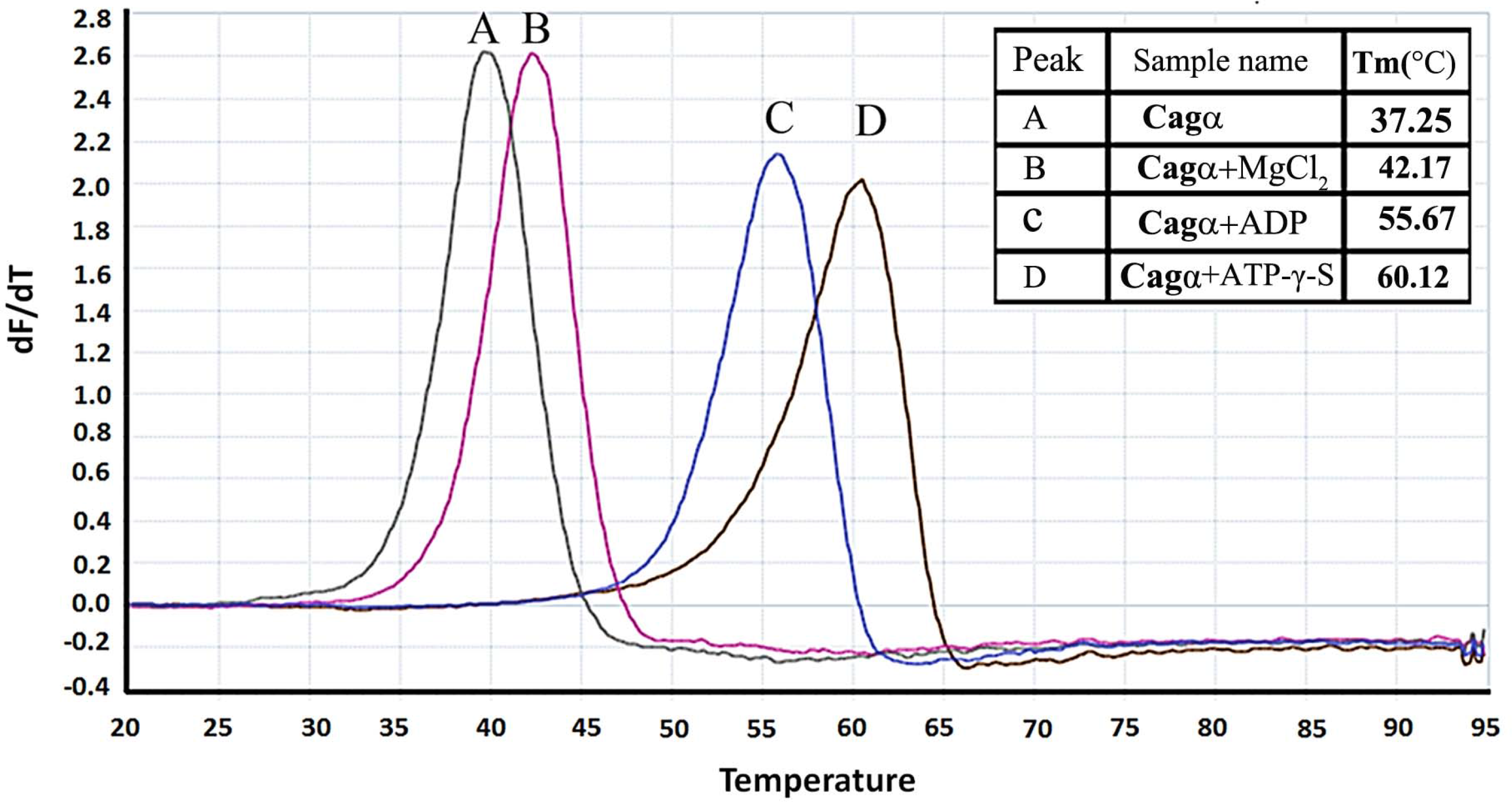
Melting temperature of Cagα in the presence of ligands and cofactors. Melting curves for Cagα were determined using differential scanning fluorimetry (DSF). (A) Cagα apoprotein (green), (B) Cagα and metal cofactor MgCl_2_ (pink), (C) Cagα with ADP (blue) and (D) Cagα with ATP-γ-S (black). Table in the upper right corner shows melting temperatures.

### Effects of binding fragments on the Cagα ATPase activity

We used a Malachite green assay to measure the release of inorganic phosphate from ATP to assess whether the 16 binding fragments impact the enzymatic activity of Cagα. Four of the molecules reduce the ATPase activity and the IC_50_ values range between 196 μM for molecule 1G2 and 4.77 mM in case of molecule 2A5 (Table 1 and supplementary Fig. 4). We used the most potent molecule 1G2 as starting point for a limited structure-activity relationship analysis using six commercially available analogs (Table 2). Two of these molecules (1G2#5 and 1G2#6) do not inhibit the ATPase activity, three of them have higher IC_50_ values than 1G2 (1G2#1, #2 and #3), but molecule 1G2#4 has a lower IC_50_ value of 81.9 μM (Table 2 and supplementary Fig. 5). Finally, we tested the mechanism of inhibition by varying the inhibitor concentrations (0 to 500 μM) and the ATP concentrations (0 to 80 μM) and fitting of the initial velocity data using nonlinear regression shows that only the V_max_ was affected, whereas the *K*_m_-values remain constant (Fig. 2). These results suggest that both molecules are non-competitive inhibitors of Cagα.

**Figure 2.**
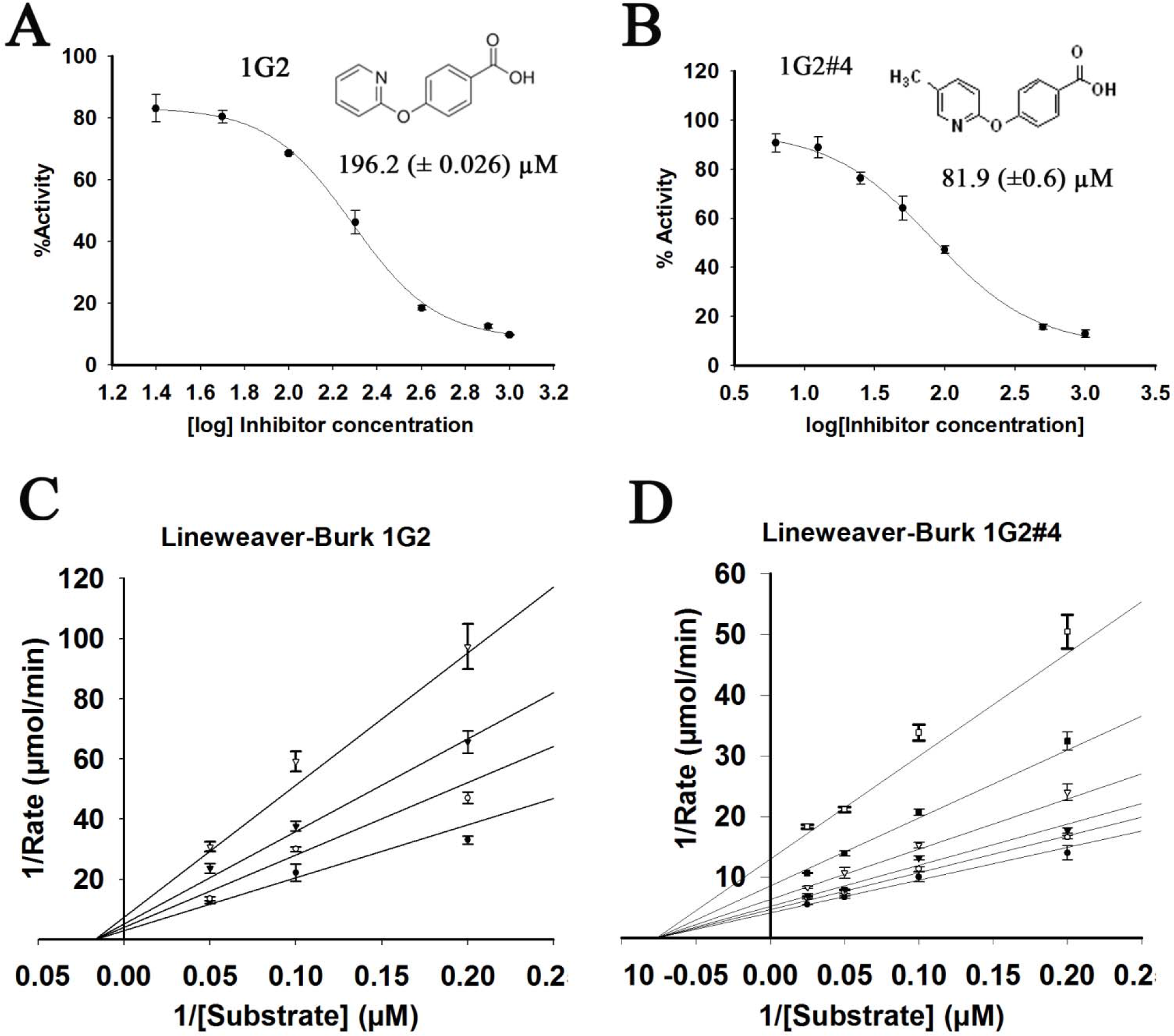
Enzyme Kinetics of Cagα in the presence of molecule 1G2 and 1G2#4. (a, c) Dose response curves of ATPase activity showing IC_50_ values in the presence of 1G2 and its derivative 1G2#4. (b, d) Lineweaver-Burke plot of Cagα ATPase activity in the presence of 1G2 and 1G2#4. The data were globally fit to a model of non-competitive inhibition. Concentrations varied from 0 to 500 μM of inhibitors in the presence of 2 mM of MgCl_2_.

**Table 1:**
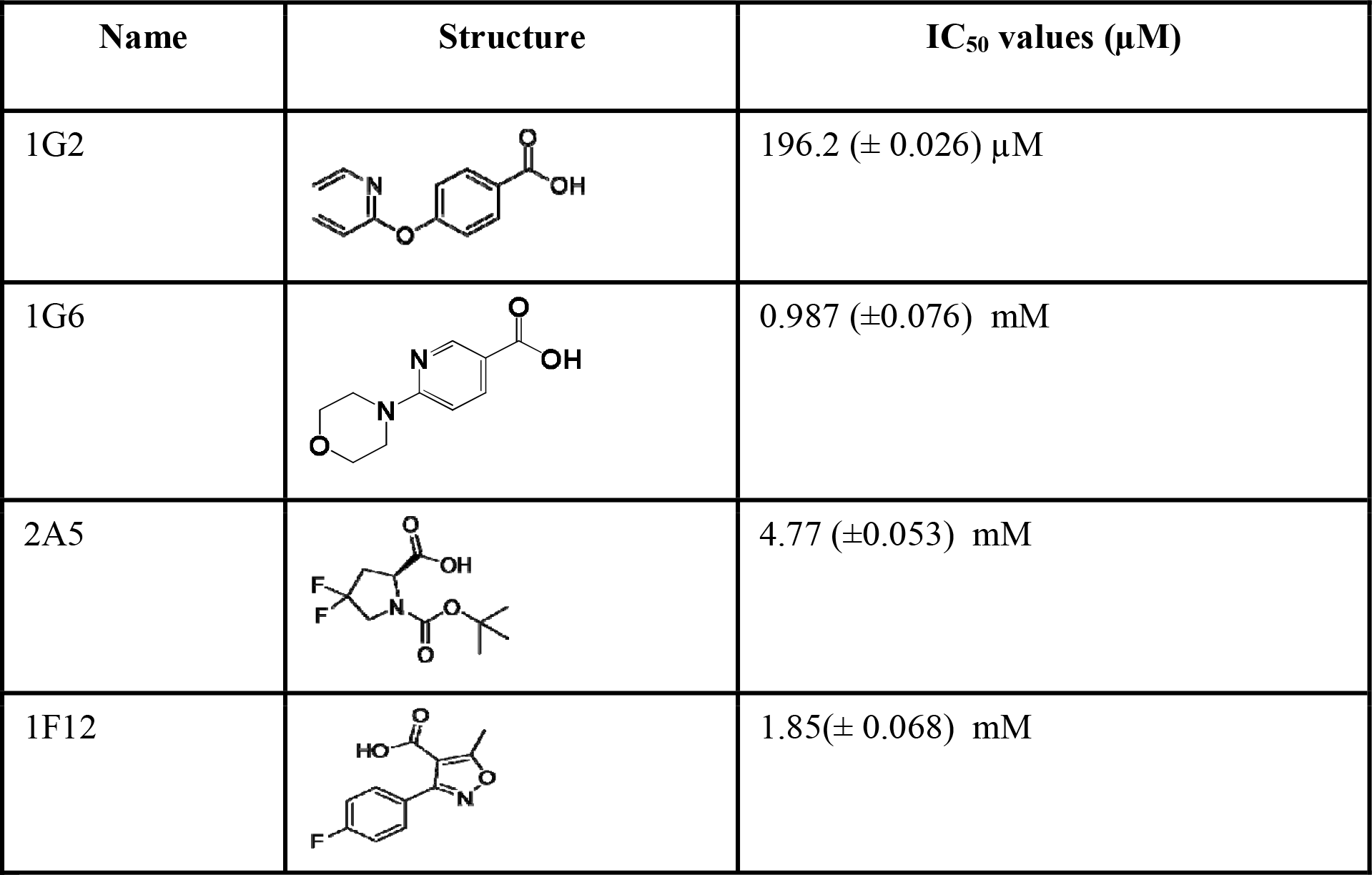
Structures and IC_50_ of molecules that inhibit the ATPase activity of Cagα.

| Name | Structure | IC<sub>50</sub> values (μM) |
| --- | --- | --- |
| 1G2 | | 196.2 (± 0.026) μM |
| 1G6 | | 0.987 (±0.076) mM |
| 2A5 | | 4.77 (±0.053) mM |
| 1F12 | | 1.85(± 0.068) mM |

**Table 2:**
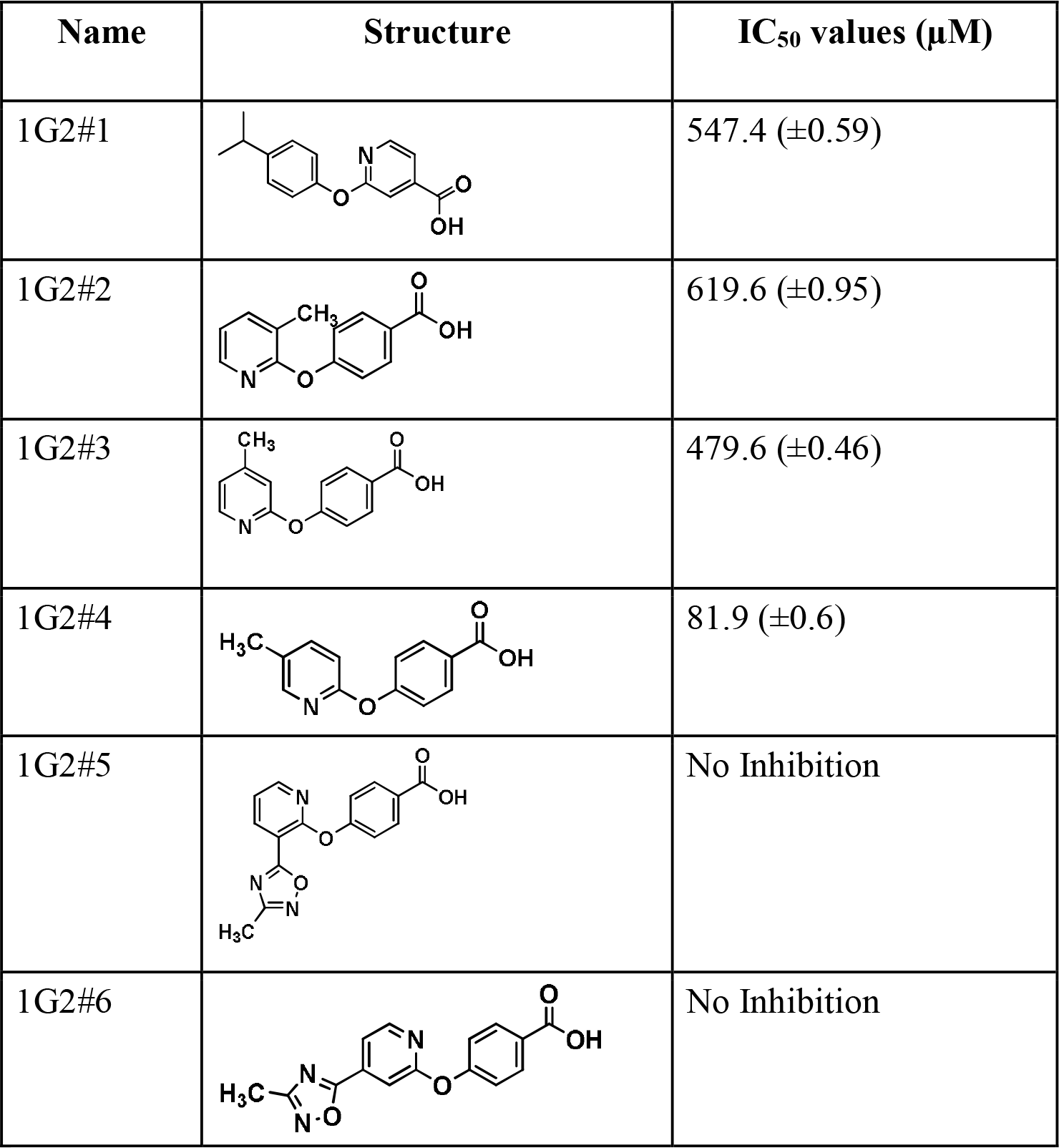
Structures of derivates of molecules 1G2 and their effects on the ATPase activity of Cagα.

| Name | Structure | IC<sub>50</sub> values (μM) |
| --- | --- | --- |
| 1G2#1 |  | 547.4 (±0.59) |
| 1G2#2 |  | 619.6 (±0.95) |
| 1G2#3 |  | 479.6 (±0.46) |
| 1G2#4 |  | 81.9 (±0.6) |
| 1G2#5 |  | No Inhibition |
| 1G2#6 |  | No Inhibition |

### Binding fragments impact the conformation and dissociate the Cagα hexamer

Binding of fragments may impact the conformation and the homo-multimerization of Cagα and we used the homo-bifunctional cross-linking agent disuccinimidyl-suberate (DSS) to obtain insights into the multimerization of the protein. As expected, incubation of Cagα with increasing concentrations of DSS (0–20 μM), followed by SDS-PAGE and western blot analysis, leads to the successive formation of higher molecular mass forms, which is consistent with the formation of a hexamer (Fig. 3a). The cross-linking pattern is similar in the presence of MgCl_2_ (Fig. 3b), ADP/MgCl_2_ (Fig. 3c), and ATP-γ-S/MgCl_2_ (Fig. 3d), but in the presence of molecule 1G2 (Fig. 3e) a reduced amount of higher molecular mass complexes is observed suggesting significant changes of the conformation and/or multimerization.

**Figure 3.**
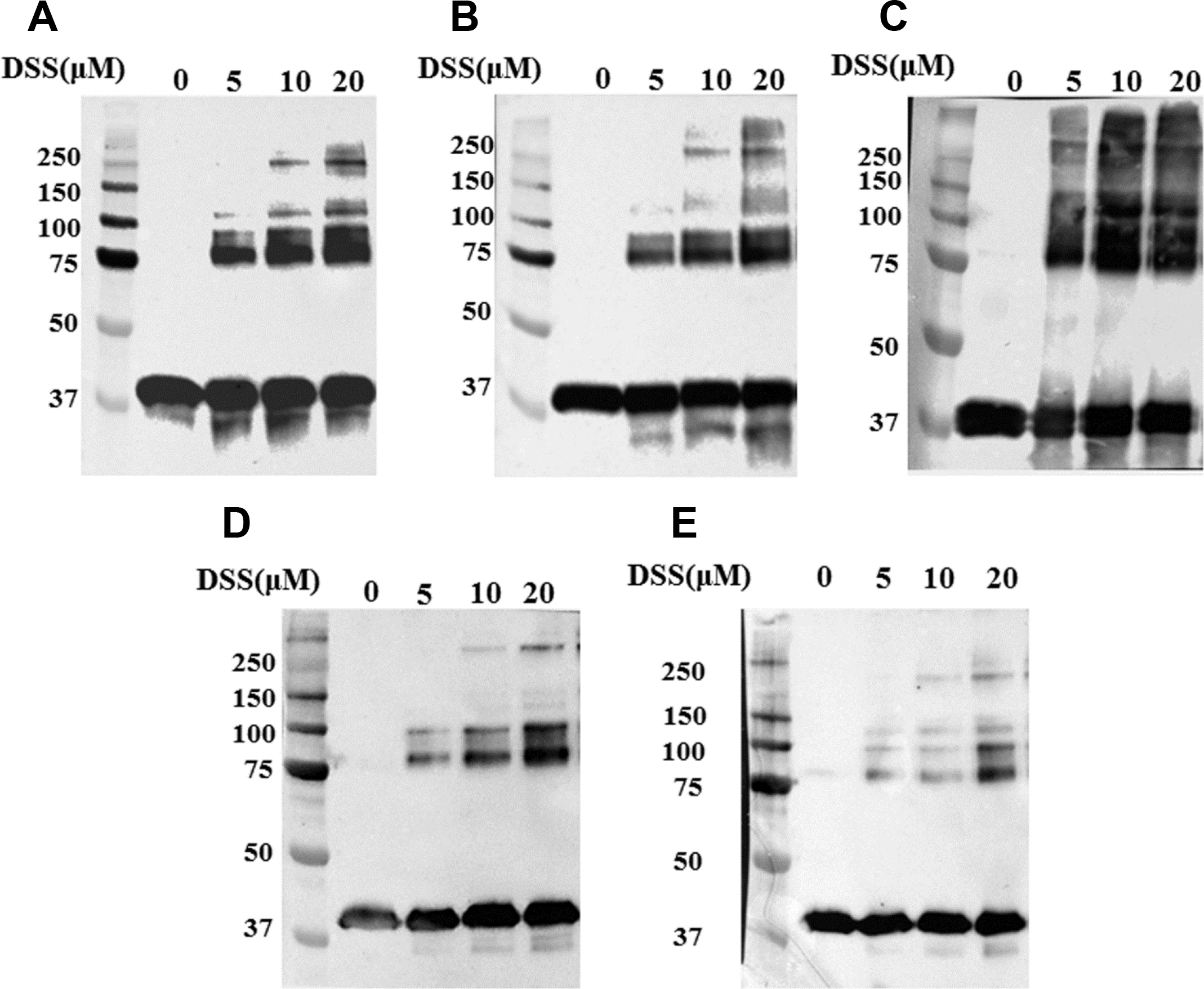
Chemical cross-linking using DSS to study the formation of Cagα oligomers in the presence of ligands. a) Cagα apo protein; b) Cagα with MgCl_2_; c) Cagα with ADP and MgCl_2_; d) Cagα with ATP-γ-S and MgCl_2_; e) Cagα with 1G2 and MgCl_2_. The concentrations of DSS varied between 0 and 50 μM leading to formation of oligomers (indicated by arrows), detection by SDS-PAGE and western blotting using His-tag specific antibodies.

To directly address this question, we performed analytical gel filtration. Purified Cagα elutes in a single peak with an elution volume corresponding to a molecular mass of 244 kDa, which is consistent with the formation of a hexamer (Fig. 4). The same elution volume was observed in the presence of ATP-γ-S, but interestingly, when Cagα was pre-incubated with molecule 1G2 we observe the elution of two peaks (peak A and peak B in Fig. 4) with elution volumes corresponding to apparent molecular masses of 175 kDa and 54 kDa, respectively. These results suggest that incubation with molecule 1G2 dissociates the Cagα hexamer into lower molecular mass species. Analysis by negative staining electron microscopy reveals hexamers in the absence of 1G2 (Fig. 5a), lower molecule mass species in peak A (Fig. 5b) and even smaller species in peak B (Fig. 5c) confirming this interpretation.

**Figure 4.**
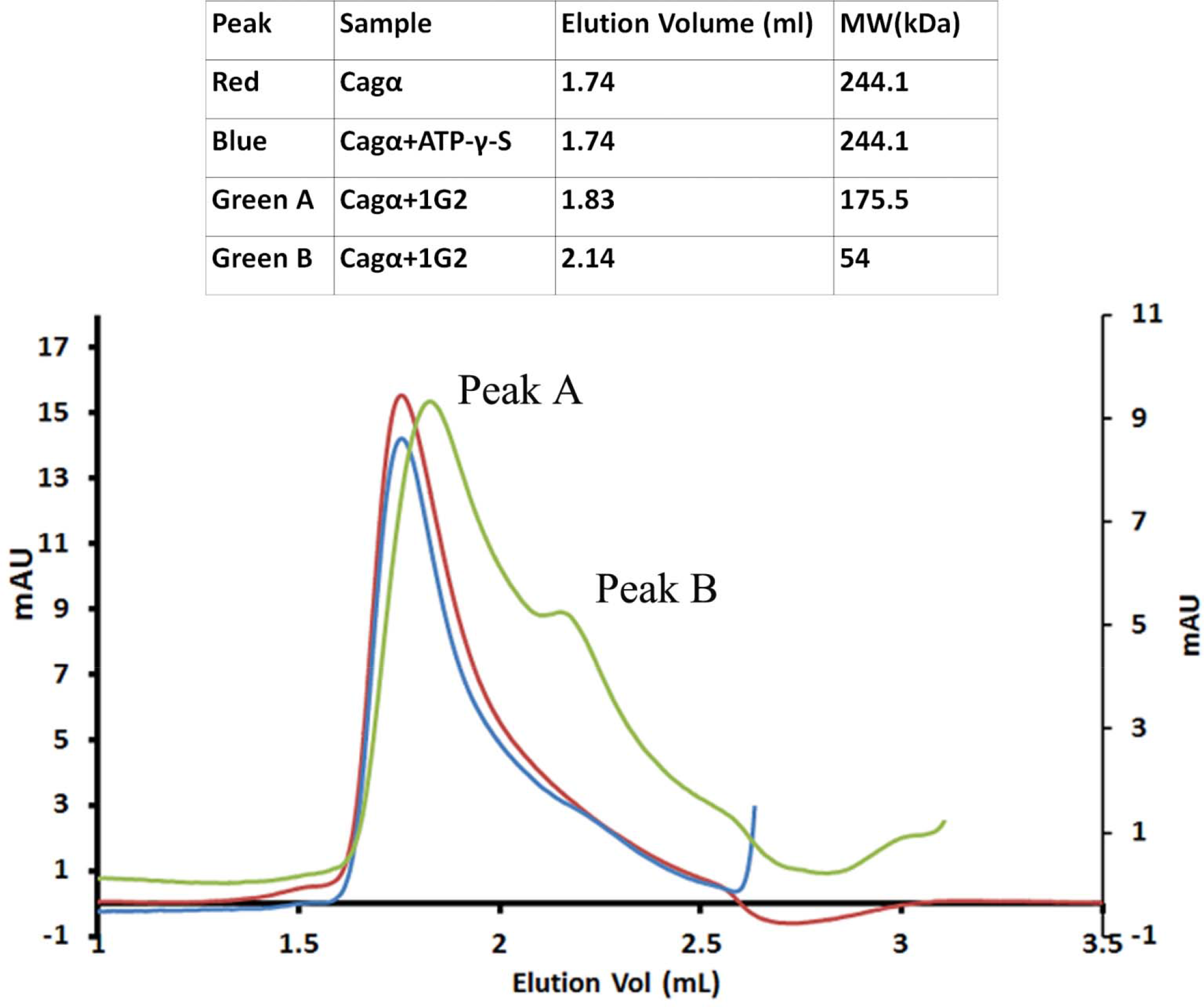
Analytical size exclusion chromatography of Cagα apoprotein and in the presence of ligands. Proteins were separated by gel filtration over a Superdex 200 column. Cagα apoprotein elutes as a hexamer (red curve), elution of Cagα-ATP-γ-S (blue curve) and of two lower molecular mass peaks (A and B) after incubation of Cagα with 1G2 (in green). The molecular masses characterized according to the elution volume are summarized in the table above the graph.

**Figure 5.**
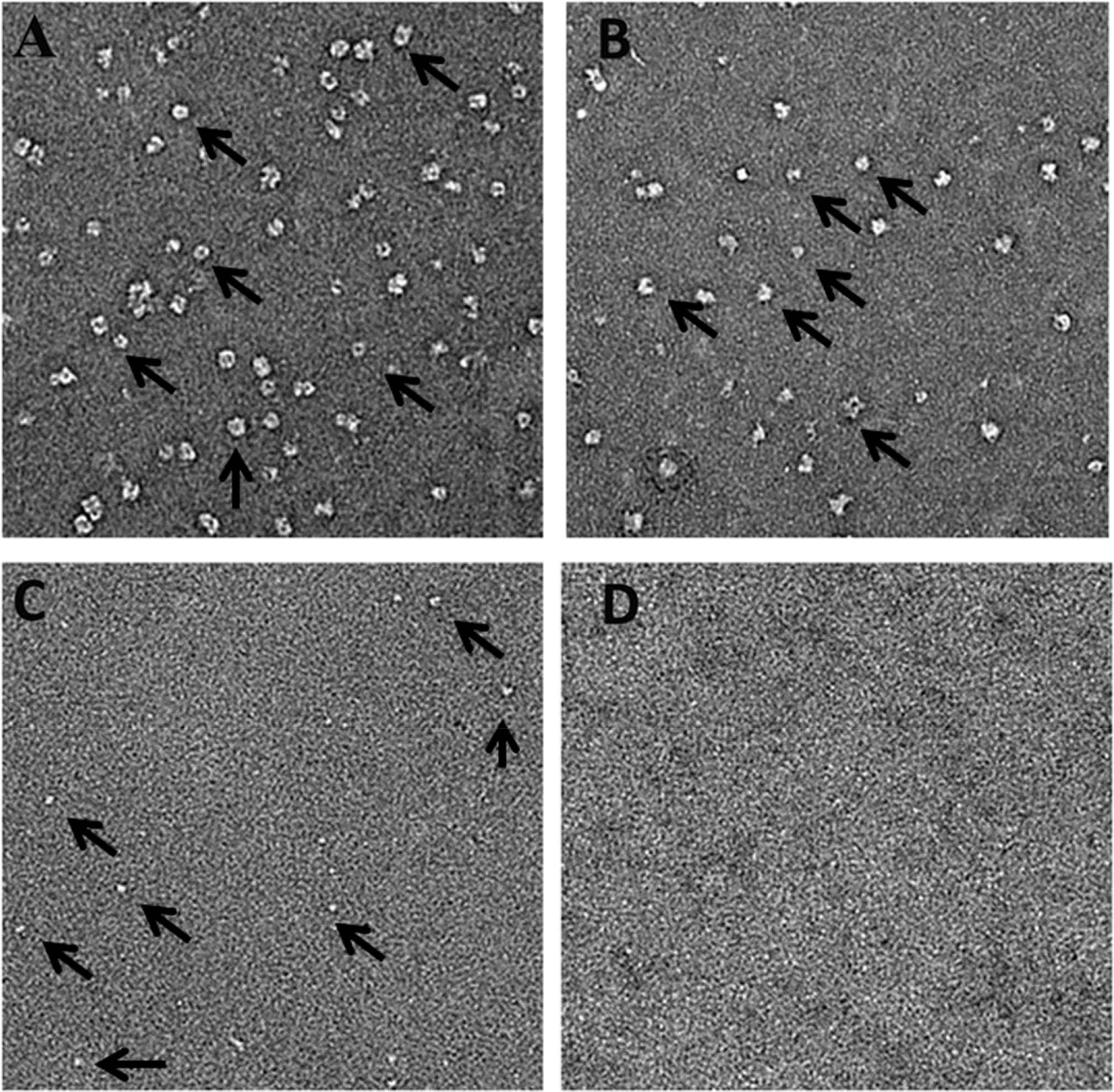
Electron micrographs of negatively stained Cagα apoprotein after gel filtration. Analysis by transmission electron microscopy and negative staining of a) Cagα apoprotein shows a hexameric ring-like structure; b) peak A of Cagα incubated with 1G2 after elution from the gel-filtration and c) peak B. d) negative control grid. Arrows show the differently sized complexes.

### X-ray analysis reveals the 1G2 binding site and conformational changes

To gain further insights into the mechanism of inhibition, we solved the X-ray structure of the Cagα-1G2 complex in the P6322 space group with two molecules in the asymmetric unit (Fig. 6a) to a resolution of 2.9 Å (Table 3). The structure was solved by molecular replacement using ADP bound Cagα (PDB code: 1G6O) as a search model. The overall structure of 1G2-bound Cagα is similar to that of the Cagα-ADP complex, but there are differences of interactions at the protein interface and we identified the electron density of molecule 1G2 sandwiched between two Cagα molecules (Fig. 6a). The monomer structure of the Cagα-1G2 complex displays both NTD and CTD with nine α-helices labeled as α1 to α9 and 13 β-strands labeled as β1 to β13 (Fig. 6b). A structural overview from the top of the NTD reveals that 1G2 interacts with the NTD of both protein subunits (Fig. 6c). The 1G2 binding site is distinct form the active site to which ADP and the substrate analog ATP-γ-S bind [16, 17]. Molecule 1G2 binds to a hydrophobic pocket created by the interaction between the NTDs of two Cagα subunits and amino acids F68 and F39 make hydrophobic contacts with the two phenyl rings of the inhibitor. R73 and D69 are the amino acids involved in forming a polar contact with 1G2. R73 interacts with the pyridine group of the second phenyl ring via a hydrogen bond and the carboxylic group of 1G2 interacts with the backbone NH group of D69 forming a potential hydrogen bond (Fig. 6d). Structural alignment of the Cagα-1G2 complex with Cagα-ADP (PDB code: 1G6O) shows an overall similar structure (RMSD 0.6 Å), but we observe shifts of the β6 and β7 sheets and in the linker region between NTD and CTD (supplementary Fig. 6). Alignment of Cagα-1G2 with Cagα apoprotein (PDB code: 1NLZ) reveals slight conformational differences in both the CTD and NTD (RMSD 0.9 Å). The α8 and α9 helical region of the CTD as well as the α1 region of the NTD display changes showing that binding to molecule 1G2 impacts the conformation of the protein (supplementary Fig. 6).

**Figure 6.**
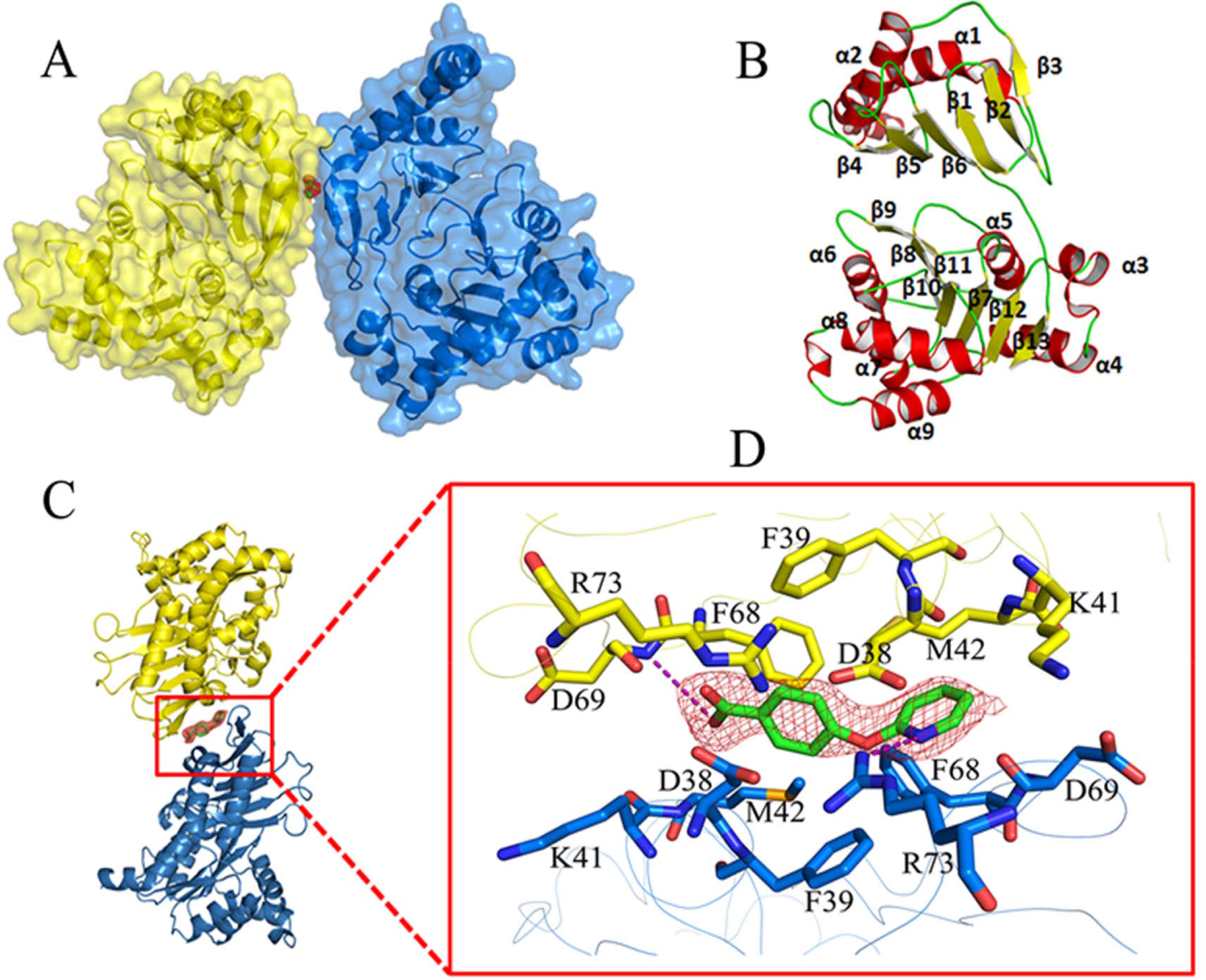
Crystal structure of the Cagα bound to molecule 1G2. a) Cartoon representation of the crystal structure of Cagα crystallized as two molecules in the asymmetric unit. Red map in middle of two subunits represents molecule 1G2. b) Representation of the monomeric subunit of Cagα in ribbon form: α helices, β strands and loops are represented in yellow, red and green, respectively. The nine helices are labeled as α1 to α9 and the β-strands are labeled as β1 to β13. c) Side view of the interaction of two subunits of protein with 1G2 in the middle represented as green stick and red map. d) Enlarged view of 1G2 binding at the interface between two protein subunits. The 2F_O_-F_C_ electron density map of 1G2 was contoured at 1.5σ.

**Table 3:**
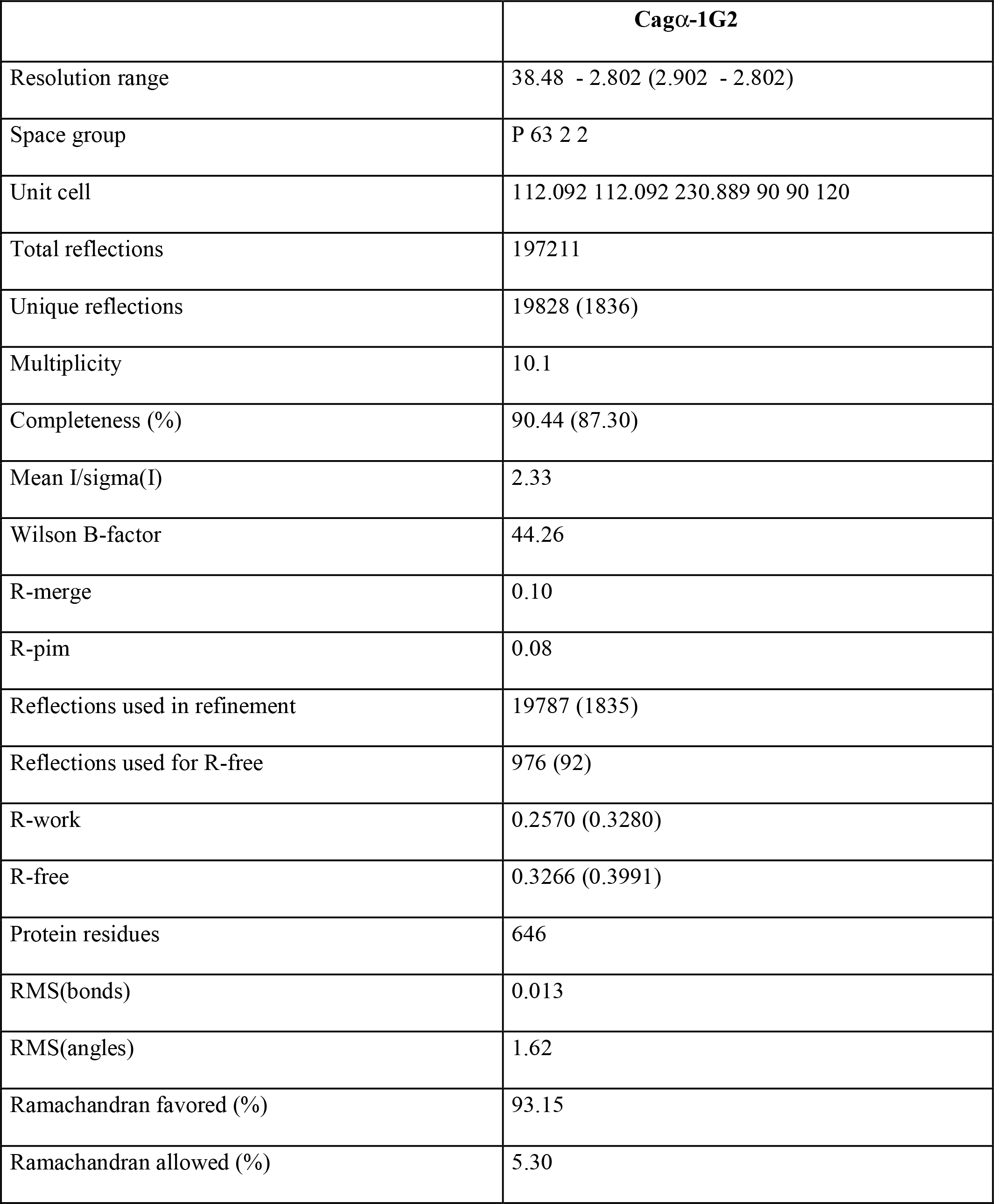
Data collection and refinement statistics.

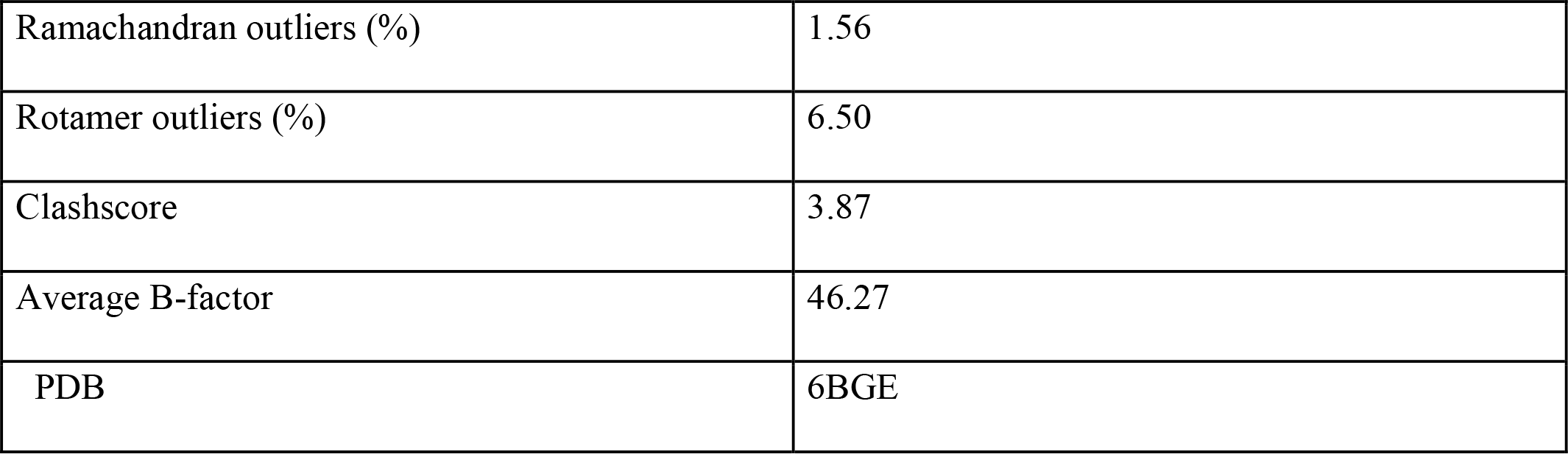

### Molecule 1G2 inhibits the production of interleukin-8 upon binding of *H. pylori* to AGS cells

Finally, we assessed whether molecule 1G2 or its derivates impact the functionality of the T4SS *in vivo*. To this effect, we tested their impact on the interaction of *H. pylori* strain 26695 with gastric adenocarcinoma (AGS) cells. First, we tested their toxicity and found that molecule 1G2 and derivates 1G2#1 to #6 have no negative effect on the growth of *H. pylori* on solid agar media at concentrations up to 500 μM (supplementary Fig. 7). Similarly, most molecules do not have negative impact on the viability of AGS cells at concentrations up to 500 μM, showing that they are not toxic (supplementary Fig. 8). When we tested the effects of these molecules at 200 μM concentration on the production of IL-8 produced by AGS cells upon co-cultivation with *H. pylori*, 1G2 significantly reduces the production of this proinflammatory cytokinin to about 50% of the control (Fig. 7). In contrast, derivates 1G2#1 to #6 have no effect on IL-8 production and none of the molecules reduces the tyrosine phosphorylation of translocated CagA, which is generally used as an alternative assay to measure T4SS function (supplementary Fig. 9).

**Figure 7.**
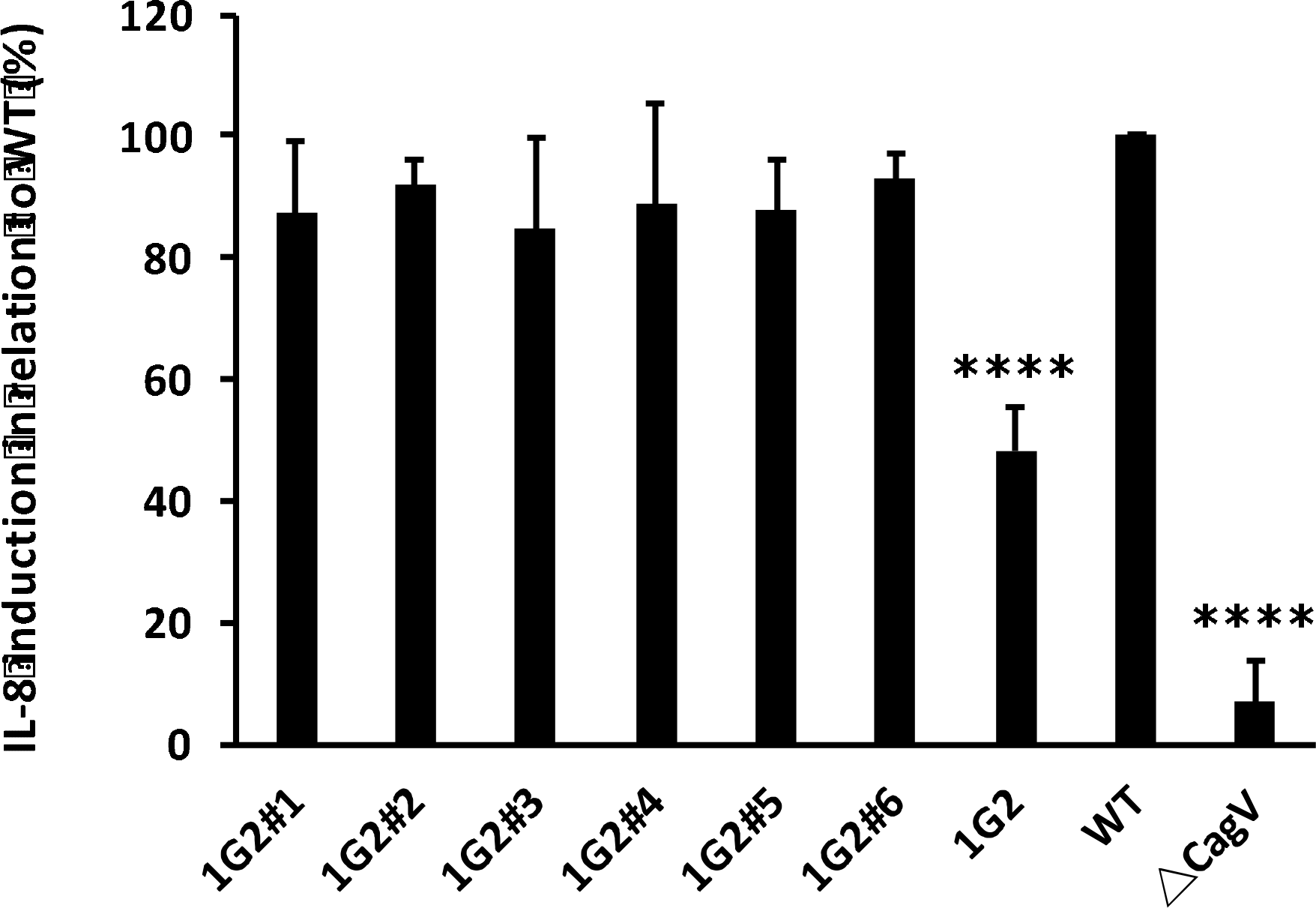
Molecule 1G2 decreases IL-8 induction in co-cultivated AGS cells. *H. pylori* 26695 without and after pre-incubation with 1G2 and its derivatives for 40 min. AGS cells were then cocultured with *H. pylori* overnight and IL-8 induction was measured by ELISA. The induction of IL-8 by the wild type was calculated as 100% (WT), induction of IL-8 by the *ΔcagV* strain was used as negative control. The data represent the results from three experiments.

## Discussion

The fragment-based screening strategy presented here was developed to identify inhibitors of protein-protein interactions and it is generally applicable to bacterial virulence factors for which structural information is available. We have previously used this approach to identify inhibitors of the VirB8 homolog TraE from the plasmid pKM101 conjugation system. We identified molecules that target a known inhibitor binding site on VirB8-like proteins, and we also identified a new binding site showing the potential for the discovery of bioactive molecules and of novel inhibitor target sites [23, 24, 33]. We here used a similar unbiased DSF-based screen to identify fragments that bind Cagα without specifically targeting its ATPase active site. Competitive inhibitors of Cagα ATPase activity are already available [29, 30], and whereas these molecules may have potential for development into antivirulence drugs, the possibility that they bind other ATPases in bacteria or in host cells remains a concern.

We here identified molecules that inhibit the Cagα ATPase activity indirectly via a novel allosteric mechanism. Enzyme kinetic analyses showed that the mechanism of inhibition by molecules 1G2 and 1G2#4 is non-competitive. Analysis of the X-ray structure showed that molecule 1G2 binds at the interface between Cagα molecules that multimerize via the NTD and this is consistent with the results of enzyme kinetics. The non-competitive mechanism of inhibition is similar to that reported in the case of unsaturated 2-alkynoic fatty acids that inhibit the VirB11 homolog TrwD from plasmid R388 [27, 28]. Docking predicted a potential binding site for these molecules at the linker region between the NTD and the CTD of a structural model of TrwD. However, high-resolution structural information on TrwD and on the potential binding site is not available and this site is distinct from the 1G2 binding site we observe by X-ray crystallography. Analysis of the Cagα-1G2 X-ray structure revealed subtle conformational changes in different parts of the protein including the active site as compared to the apoprotein and its complex with ADP. This may explain the effect of 1G2 binding on the enzymatic activity of Cagα. Conformational changes were also observed by cross-linking and comparable observations were made in the case of TrwD that became more susceptible to protease degradation in the presence of 2-alkynoic fatty acids [27]. Intriguingly, gel filtration and EM analysis revealed that binding to 1G2 successively dissociates the Cagα-hexamer. This mechanism is entirely novel and it would be interesting to assess the molecular basis of dissociation using approaches that are more sensitive to conformational changes than X-ray crystallography, such as cryo-electron microscopy. Also, in the context of the current work we have not assessed the potential of most of the other Cagα-binding molecules that do not inhibit the ATPase activity. These molecules may bind different sites of the protein and may have interesting biological activity that remains to be explored in future.

Cagα is essential for type IV secretion and deletion of the encoding gene inhibits IL8 production and Cagα transfer into AGS cells, which are the most commonly used readouts for *H. pylori* T4SS function [9, 10]. The results for IL-8 production and CagA phosphorylation usually correlate when the effects of *cag* gene mutations are determined. It was therefore somewhat unexpected that molecule 1G2 inhibited IL-8 production to 50% of the control values, but we did not observe an effect on CagA phosphorylation. This observation may be due to the partial inhibition of T4SS function by 1G2 that is more readily quantifiable in the IL-8 production assay as compared to the CagA phosphorylation assay. Enzyme kinetic analysis showed that molecule 1G2#4 was significantly more potent than 1G2 *in vitro*, but repeated crystallisation trials of Cagα with 1G2#4 were unsuccessful. Soaking of 1G2#4 into Cagα crystals resulted in crystal cracking, suggesting a conformational change in Cagα and/or dissociation of the Cagα quaternary structure. This may be due to the higher potency of molecule 1G2#4 that could be explained by additional hydrophobic contacts of its additional methyl group with amino acid K41 (supplementary Fig. 10). However, this molecule had no effect in the *in vivo* assays, which may be due to its higher hydrophobicity impacting solubility and penetration into cells.

We have here analyzed six derivates of molecule 1G2 that were commercially available and in future work we will conduct a structure-based structure-activity relationship analysis to synthesize more potent molecules that efficiently penetrate into cells. This approach will enable us to assess whether inhibition of Cagα leads to differential effects on the translocation of effectors CagA, HBP and murein that might also explain the differential effect of molecule 1G2 on IL-8 production and CagA phosphorylation. Potent inhibitors of Cagα could be developed into anti-virulence drugs that are alternative or complementary treatments to currently used triple or quadruple therapy. It would also be interesting to test the specificity of these molecules to assess whether they are narrow or broad spectrum inhibitors that also impact other T4SS, e.g. bacterial conjugation systems [28, 34].

## Methods

### Bacterial strains, cell lines and culture conditions

*H. pylori* strains 26695 and *ΔcagV* (*hp0530*) mutant have been described [35] and were cultivated on Columbia agar base (BD) containing 10% (v/v) defibrinated horse blood (Winsent Inc.), vancomycin (10 μg/L) and amphotericin B (10 μg/L). Chloramphenicol (34 μg/L) was added in case of the *ΔcagV* strain to select for the cam gene cassette used to disrupt the gene. For liquid culture, brain heart infusion (BHI) media (Oxoid) were supplemented with 8% fetal bovine serum (FBS) and appropriate antibiotics. Bacteria were cultivated at 37□, under microaerophilic conditions (5% oxygen, 10% CO_2_). AGS cells were grown at 37□ in F12K media (Winsent Inc.) with 10% (v/v) FBS (Winsent Inc.) in a 5% CO_2_ containing atmosphere.

### Cloning, expression and purification of Cagα

The Cagα encoding gene from *H. pylori* 26695 (ATTC) was PCR-amplified from genomic DNA with primers (forward, 5’-TAGCGAATTCGGTACCATGACTGAAGACagαTTGAGTGCA-3’ and reverse, 5’-CGATGAATTCCTCGAGCTACCTGTGTGTTTGATATAAAATTC-3’), thereby adding an N-terminal hexahistidyl-tag and a TEV cleavage site, and the PCR product was cleaved with restriction enzymes *NheI* and *XhoI*, followed by ligation into expression vector pET28a. Expression was conducted in *E. coli*BL21 (DE3) cultivated in two liters of LB-medium at 37 °C at 220 rpm, protein production was induced at OD_600_ of 1 with 1 mM isopropylthio-β-galactoside (IPTG), followed by further incubation for 16h at 25°C. For purification, the cell pellet was suspended in binding buffer (50 mM HEPES, 500 mM NaCl, 20 mM imidazole, pH 7.5, 10% glycerol, 0.1% triton, plus two tablets of EDTA-free protease inhibitor cocktail (Roche)) and lysed using a cell disrupter (Constant Systems Inc.) at 27 kPsi, followed by centrifugation at 15,000 rpm at 4°C to reduce cell debris. The supernatant was loaded onto a His-trap Ni-NTA column (GE Healthcare), which was pre-equilibrated with 100 ml of binding buffer. Then protein was eluted using a linear 50 ml gradient of 40-500 mM imidazole in binding buffer. Proteins were then desalted into TEV buffer (25 mM sodium phosphate, 125 mM NaCl, 5 mM DTT, pH 7.4) and subjected to cleavage of the N-terminal 6x-His-tag using TEV protease in a ratio of 1:70 (TEV:protein) for 24 h at 20°C. The cleaved protein was dialysed into 50 mM HEPES, 100 mM pH 7.5 buffer and concentrated with Amicon filters. Size exclusion chromatography was conducted using a Superdex-200 column (GE Healthcare) with buffer 25 mM HEPES pH7.5 and 100 mM NaCl and peak fractions were analyzed by SDS-PAGE. The fractions containing Cagα hexamers were pooled and concentrated to 6 mg/ml for crystallographic studies.

### Analytical gel filtration chromatography

Purified protein was further characterized by analytical gel filtration (Superdex 200) in 25 mM HEPES, pH 7.5 and 50 mM NaCl (pH 7.5). The column volume was 3ml and the protein was injected at a flow rate of 0.5 ml/min. To study the effects of ATP-γ-S and of 1G2, 35 μg of Cagα was preincubated with 2 mM of the molecules for 30 min, followed by analytical size exclusion analysis.

### Enzyme activity assay

The ATPase activity was quantified using a malachite green binding assay [36]. The 100 μL reaction mixtures contained 25 mM HEPES (pH 7.5), 100 mM NaCl, 60 nM of enzyme and 200 μM of MgCl_2_ with different concentrations of ATP (0μM - 320μM) to determine kinetic parameters. The reaction mixtures were incubated for 30 min at 30°C and then 40 μL of malachite green assay mixture was added. The formation of the blue phosphomolybdate-malachite green complex was in linear relation to the amount of released inorganic phosphate and measured at 610 nm. To study the mechanism of inhibition, the concentrations of inhibitors were varied between 0 and 500 μL with different concentrations of ATP (0-40 μM). Initial velocity data were fit using nonlinear regression analysis to each of the equations describing partial and full models of competitive, uncompetitive, noncompetitive, and mixed inhibition using the Enzyme Kinetics Module of SigmaPlot (SigmaPlot version 11.0 software). On the basis of the analysis of fits through “goodness-of-fit” statistics, the full noncompetitive inhibition model was determined with the equation v = *V*_max_/[(1 + [I]/*K*_i_) × (1 + *K*_m_/[S])], where [S] = [ATP], [I] = [1G2].

### IC_50_ determination

IC_50_ values were determined by incubating different concentrations of molecules (10 - 1,000 μM; from stocks of 200 mM) with enzyme in 25 mM HEPES (pH 7.5) and 100 mM NaCl. Mixtures were incubated with inhibitors for 15 min, followed by addition of ATP and incubation for 30 min at 37 °C. The reactions were stopped by addition of 40 μl malachite green solution and the inorganic phosphate released was determined at 610 nm. Data were plotted as 1/rate versus inhibitor concentration for each substrate concentration and a linear fit was calculated by non-linear regression using SigmaPlot (version 11.0).

### Differential scanning fluorimetry (DSF)

A fragment library of 505 molecules was used as in our previous work [32]. The reaction mixture contains 5μM of Cagα, 10x concentration of SYPRO Orange (from 5000x stock solution (ThermoFisher)) in 50 mM HEPES (pH 7.5), 100 mM NaCl and 5% final concentration of DMSO. The fragments and nucleotides were added to final concentrations of 5 mM, and the fluorescence was monitored over 20-95 °C with a LightCycler 480 instrument (Roche).

### Crystallisation and structure determination

Initial crystallization conditions were established using the MCSG screen from (ANATRACE, USA) using 6 mg/ml of Cagα and 1 mM of 1G2 (1:10 ratio). Final crystals were grown at room temperature using the hanging drop vapour diffusion method in 100 mM Bis-Tris (pH 6.5) and 2 M ammonium sulfate. Drops containing 2 μl of protein-inhibitor-mixture (1:10 ratio) and 2 μl of reservoir solution were incubated for 2 weeks Hexagonally-shaped crystals appeared after 7-10 days. The crystals were cryo-protected in 100 mM Tris-HCl buffer (pH 8.5), 2 M ammonium sulfate and 25% glycerol, flash frozen in liquid nitrogen and the data were collected at microfocus beamline F1 at the Cornell High Energy Synchrotron Source (CHESS). The intensity data was processed using the HKL2000 [37] program in p6522 space group (Table 3) . The structure was solved by molecular replacement using the coordinates of PDB ID: 1G6O as search model. Refinement and modeling were performed using REFMAC and Coot [38, 39]. Final graphical figures and tables were generated using the Pymol-integrated Phenix software suite [40].

### Analysis of protein-protein interactions by cross-linking

Chemical cross-linking with disuccinimidyl suberate (DSS; Pierce) was performed as described [41]. 100 nM of Cagα in 50 mM HEPES (pH 7.5) and 100 mM NaCl were first incubated with cofactors (MgCl_2_, ADP) or inhibitors (ATP-γ-S, 1G2) for 30 min, followed by crosslinking with DSS (0 - 50 μM) for 1 h, and reactions were stopped by mixing with an equal volume of 2 × Laemmli buffer. The formation of cross-linking products was analyzed by SDS-PAGE and western blotting using His-tag specific antiserum and ImageLab 4.0 software (Bio-Rad)

### Electron microscopy and image processing

Carbon-coated grids were negatively glow-discharged at 15 mA and 0.4 mBar for 30 sec. 5 μl of purified protein at a concentration of 2 ng/μl was spotted onto the grids for 60 sec and blotted using grade 1 Whatman filter paper, followed by staining with freshly prepared 1.5% uranyl formate solution for 60 sec and drying. The samples were imaged at a magnification of 49,000-fold (pixel size: 2.2 Å/pixel) with a defocus of −2.5 μm using a FEI Tecnai T12 electron microscope (FEMR facility at McGill University). Transmission Electron Microscope (TEM) equipped with a Tungsten filament and operated at 120 kV equipped with a 4k × 4k CCD camera (Gatan Ultrascan 4000 CCD camera system model 895). Subsequently, the images were processed using ImageJ.

### Measurement of *H. pylori* and AGS cell viability

AGS cell viability was monitored using Cell Proliferation Reagent WST-1 (Sigma). To evaluate the sensitivity of *H. pylori* to 1G2 and its derivates, freshly harvested bacteria were spread on a 150mm agar plate. Increasing concentrations of compounds (50-500 μM) were spotted onto Whatman paper disks and growth was observed after 72h incubation at 37°C under microaerophilic conditions and compared to antibiotics (50–250 μM).

### Assay for monitoring CagA transfer into AGS cells

Preceding the infection, an overnight culture of *H. pylori* was pre-incubated with 1G2 and its derivates for 30 min. AGS cells at 6 × 10^5^ cells/well density in 6-well plates were infected with the pretreated cultures of *H. pylori* for 3-6 h at a multiplicity of infection of 100:1. Cells were washed twice with PBS, harvested and lysed at 4°C in RIPA buffer (150 mM NaCl, 50 mM Tris/ HCl, pH 8, 1% NP-40, 2 mM Na_3_VO_4_, supplemented with Complete Protease Inhibitor Tablet (Roche). After 15 min of centrifugation at 16,000 *g*, lysates were separated by SDS-PAGE, followed by western blotting with mouse polyclonal antiserum raised against CagA (Abcam), anti-phosphotyrosine (PY99; Santa Cruz Biotechnology) and anti-β-actin (C4, Santa Cruz Biotechnology).

### Assay for IL-8 induction

Preceding the infection, an overnight culture of *H. pylori* was pre-incubated with 1G2 and its derivates for 30 min. AGS cells at 6 × 10^5^ cells/well density in 6-well plates were infected with the pretreated cultures of *H. pylori* at a multiplicity of infection of 100:1. After 24 h incubation under microaerophilic conditions, supernatants were sampled and centrifuged (15,000 g), before freezing at - 80°C. The level of IL-8 in cell culture supernatants was determined by using a commercially available human IL-8 ELISA kit (Invitrogen).

## Acknowledgements

This work was supported by grants to C.B. from the Canadian Institutes of Health Research (CIHR MOP-84239)(http://www.cihr-irsc.gc.ca/), the NSERC-CREATE program on the Cellular Dynamics of Macromolecular Complexes (CDMC) (http://www.nserc-crsng.gc.ca/), a seed grand from Merck, Sharp and Dohme (http://www.merck.ca/), the Canada Foundation for Innovation (CFI)(https://www.innovation.ca/) and the Fonds de recherche du Québec-Santé (FRQ-S)(http://www.frqs.gouv.qc.ca/). We are grateful to Edward Ruediger and his colleagues at the medicinal chemistry platform at IRIC (Institut de recherche en immunologie et en cancérologie (IRIC), Université de Montréal) for support with small molecule screening. We thank Dr. Aleksandr Sverzhinsky from the Department of Biochemistry and Molecular Medicine for helping us with analytical chromatography. We are thankful to Dr. Martin Schmeing and Dr. Asfarul Haque (Department of Biochemistry, McGill University) for helping us with data collection at the Facility of Electron Microscopy Research (FEMR) at McGill University. Synchrotron X-ray data were collected at the Cornell High Energy Synchrotron Source (CHESS, MacCHESS beamline F1).

